# Capturing Natural Evolution in Function-guided RNA Design via Genomic Foundation Models

**DOI:** 10.1101/2025.04.08.647793

**Authors:** Yanjie Huang, Bibi Zhang, Jack X. Chen, Runze Ma, Zhongyue Zhang, Xianjun Chen, Yi Yang, Shuangjia Zheng

## Abstract

RNAs play critical roles in gene expression regulation, catalysis, genetic information transmission, and immune response. However, current RNA engineering approaches face limitations in designing RNA for efficient in vivo folding and functional execution. Here, we present a novel, zero-shot strategy that integrates genomic large language models (LLMs) with inverse folding models (IFMs) to generate RNA sequences that mirror natural evolutionary patterns while preserving the structural integrity of critical functional regions. Benchmarking 9 state-of-the-art genomic foundation models, we show that their unsupervised log-likelihood scores correlate strongly with RNA fitness, enabling zero-shot sequence design. Applied to fluorescent RNA aptamers, our approach yielded 20 Broccoli variants, over half of which achieved up to 155% in fluorescence and a twofold increase in binding affinity, with robust performance both in vitro and in vivo. For Pepper, two rounds of optimization produced 40 variants, 16 of which demonstrated up to 2.6 fold in fluorescence and a threefold boost in binding affinity, including five with superior in vivo brightness.This approach offers a scalable, efficient, and resource-light method for RNA design, with significant implications for RNA-based therapeutics and diagnostics. By leveraging general foundation models to facilitate RNA evolution in a zero-shot fashion, we believe our work represents a major advancement in the field of RNA engineering.

## Introduction

RNA is a fundamental biomolecule whose biological functions depend on its dynamic and flexible three-dimensional conformations, enabling critical roles in essential life processes such as genetic information transmission and gene expression regulation.^1–3^ Moreover, RNA’s inherent advantages—such as transient expression in cells, efficient chemical synthesis and site-specific modification, and low immunogenicity—have spurred its increasing use in clinical therapeutics and diagnostics.^4,5^ The widespread application of RNA has driven advancements in rational design strategies, promoting the development of RNA engineering to meet diverse functional requirements. ^6–8^However, compared with naturally evolved RNA molecules refined by millions of years of evolutionary pressure in a cellular context, in vitro derived RNA molecules often result in sub-optimal folding within complex cellular environments while optimizing for target-binding capabilities, a main technical challenge particularly pronounced in RNA aptamer development. ^9^Consequently, developing RNA design methods capable of achieving efficient *in-vivo* folding and precise functional execution remains one of the core challenges in RNA engineering applications.

Currently, the SELEX (Systematic Evolution of Ligands by Exponential Enrichment) technique, as a benchmark approach for evolving and validating RNA, has been widely employed in RNA design.^10,11^ This method utilizes an *in vitro* directed evolution strategy to obtain target aptamers through multiple rounds of selection and amplification. However, SELEX not only proves time-consuming and labor-intensive but also suffers from limitations imposed by the restricted diversity of oligonucleotide libraries, thereby hindering comprehensive exploration of the entire sequence space.^12,13^ Crucially, the *in vitro* nature of SELEX selection means that it often fails to account for the complex factors that influence RNA folding in vivo. Although SELEX-selected aptamers exhibit robust target-binding capabilities under controlled conditions, the simplified selection environment does not recapitulate complex cellular context, leading to significant discrepancies in folding efficiency and ultimately limiting their potential applications in synthetic biology.^9^

With the remarkable success of general large language models (LLMs) across multiple fields, foundation models for scientific computing are emerging as a new paradigm to accelerate interdisciplinary discoveries.^14–17^ In the field of protein science, recent studies have shown that advanced foundation models can effectively guide the directed evolution of proteins, successfully designing proteins with enhanced functional properties.^16,18^ However, comparable breakthroughs remain elusive in RNA research, largerly due to two major bottlenecks. Firstly, RNA molecules exhibit dynamic structural plasticity, this inherent flexibility disrupts the establishment of reliable sequence-structure-function relationships, posing significant challenges for language models in capturing discontinuous structural features and inferring molecular functions. Secondly, while RNA sequencing data is abundant, it predominantly originates from conserved functional regions of limited species, resulting in poor functional annotation and sequence diversity; ^19,20^Moreover, the scarcity of large-scale, high-confidence structural annotations on RNA further restricts the model’s generalizability and predictive accuracy. ^21^

Here, we hypothesize that integrating a genomic large language model (LLM) capable of simulating millions of years of evolutionary variation with an inverse folding model that captures dynamic RNA structural information can generate functional RNA sequences. These sequences would align with natural evolutionary patterns at the sequence level while preserving the structural integrity of key functional regions. To validate this, we conducted comprehensive benchmarking of existing nucleotide LLMs and inverse folding models. Empirical evaluations demonstrated that both types of models can guide RNA-directed evolution and capture evolutionary patterns across distinct dimensions, enabling the construction of a multidimensional fitness landscape. Building on these insights, we propose RILLIE – “RNA In Silico Evolution via LLM and Inverse folding”, a design strategy that synergizes evolutionary constraints with structural biophysics to achieve efficient, evolutionarily informed RNA design. Using fluorescent aptamers as an example, we implement single-round and multi-round evolutionary strategies in the Broccoli and Pepper systems to address the common issue of getting trapped in local optima during directed evolution. For Broccoli, we generated 20 variants, over half of which showed significant improvements in fluorescence intensity (up to 55%). In vivo testing via living cell imaging and Fluorescence-Activated Cell Sorting (FACS) confirmed enhanced fluorescence in living cells. For Pepper, one of the best-performing fluorescent aptamers with nanomolar-level affinity, we tripled its binding affinity with only 20 sequences per design round. This iterative approach led to a 2.6 fold fluorescence intensity, with over 40% of sequences outperforming the wild-type. FACS testing also confirmed enhanced fluorescence, and the second round of directed evolution introduced 6 to 14 mutations, less than 75% similarity to the wild type while maintaining strong fluorescence.Consequently, we believe that our proposed new method holds great potential in advancing RNA evolution.

## Results

To test zero-shot performance of genomic foundation models on RNA directed evolution, we benchmarked 9 SOTA genomic foundation models. These models included RNA language models(AIDO.RNA^22^, RiNALMo^23^, RNAFM^20^, RNAMSM^24^), DNA language models (Evo^17^, Nucleotide Transformer^25^, GENA-LM^26^ and GROVER^27^) and the RNA inverse-folding model (RhoDesign^8^). We collected 6 ncRNA DMS datasets including tRNA^28^, RNA aptamer^10,11,29^ and ribozyme^30^ and evaluated the model’s ability to perform zero-shot ncRNA fitness prediction using the results of experimental ncRNA DMS studies as the ground truth score. (Materials and Methods) Notably, recent study has shown that DNA foundation models trained on large corpora of nucleotide coding sequences also exhibit comparable abilities to predict mutational effects on ncRNA function.^17^ Thus, we integrated 4 DNA language models into our benchmark. The likelihood of the nucleotide sequence, as predicted by the models, is used to estimate the experimental fitness score (Materials and Methods). We then compute the correlation coefficient between the predicted fitness score and the actual experimental score to assess the model’s performance (Fig. 1c).

**Fig. 1.**
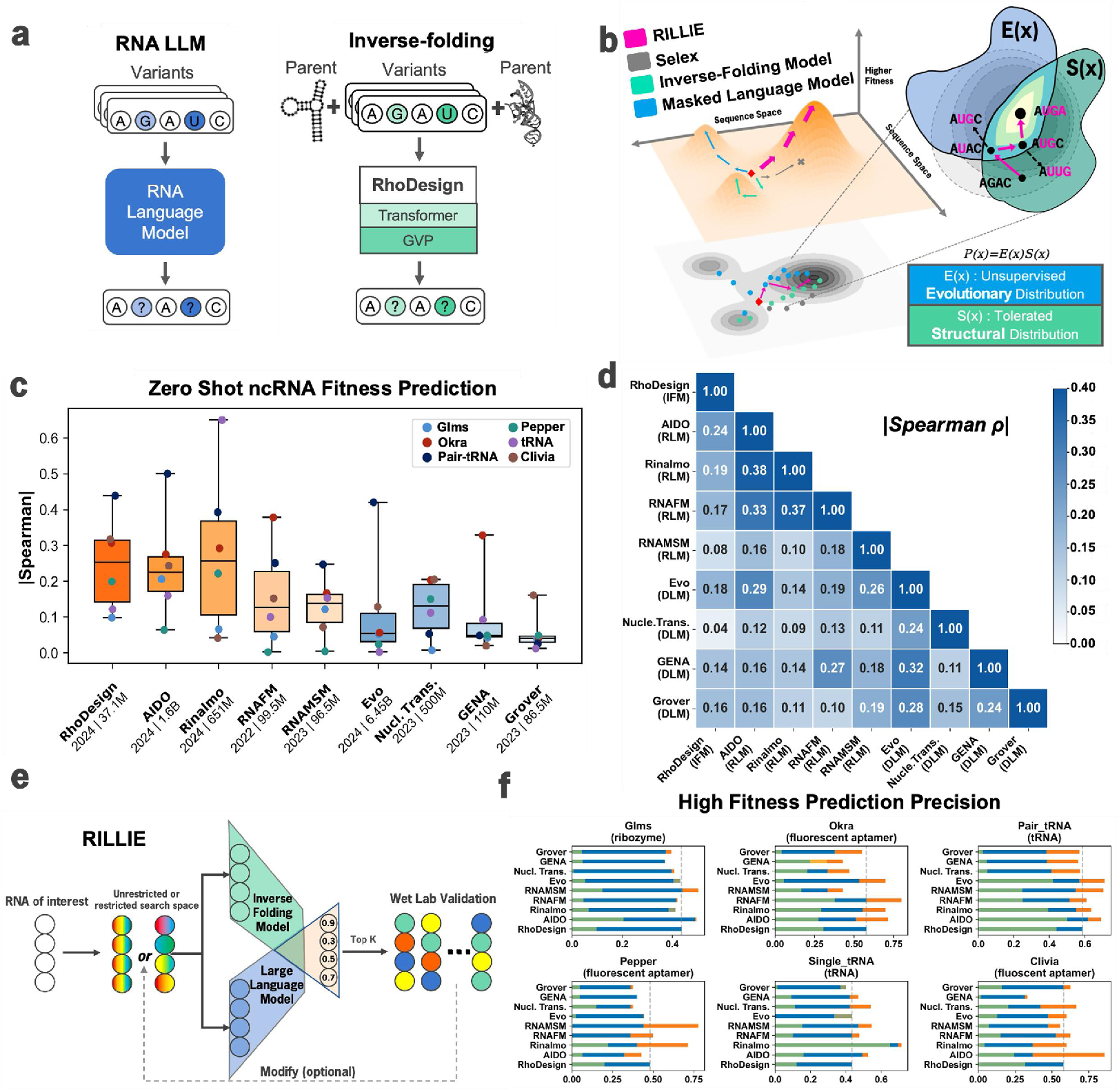
Guiding evolution of diverse RNA with both sequence and structure rules. **a**, Schematic diagram of RNA large language model (RLM) and inverse-folding model (IFM). Language models and inverse-folding models represent two different types of genomic foundational models. RNA large language models perform mask and predict learning on a large number of RNA sequences, capturing evolutionary information in the sequence dimension. In contrast, an inverse-folding model learns through masked training on structures, capturing the mapping relationship between sequence and structure as well as the structural feasibility. **b**, Schematic diagram of how combination of sequence and structure rules facilitates Directed Evolution. Theoretically, balancing sequence and structure rules can ensure that the sequence maintains structural feasibility while undergoing directed evolution, leading to a higher success rate and better properties of the variants. **c**, Zero-shot ncRNA fitness prediction. Box plots show Spearman correlations between predicted likelihoods (from various models) and measured fitness values across six DMS datasets (including fluorescent aptamers, riboswitches, and tRNAs). Red boxes indicate IFMs; orange boxes, RNA LLMs; blue boxes, DNA LLMs. Nucl.Trans. denotes Nucleotide Transformer.The results show that all foundation models have learnt certain evolutionary patterns.(detail data in each dataset in **Extended Data Fig. 1b**) **d**, The average Spearman Correlation between different models across six dataset. IF refers to inverse-folding model. RLM refers to RNA language model. DLM refers to DNA language model. The results show that different foundation models capture different evolutionary patterns. (detail data in each dataset in **Extended Data Fig. 2**) **e**, Pipeline of our model, RILLIE – “RNA In Silico Evolution via LLM and Inverse folding”, a hybrid autoregressive model integrating sequence and structure information to evaluate the joint likelihood over all positions in a sequence. RNA sequence are tokenized (green), combined with structral features extracted from a inverse-folding model (blue). The top k scoring sequences are selected for wet lab validation. Based on the experimental results, the library can be modified (if necessary). **f**, High-fitness sensitivity analysis reveals that genomic foundation models are good at picking out high fitness variants and multimodal input improves language model performance. Green bar shows Spearman Correlation in **Extended Data Fig. 1b**, blue bar shows High-fitness Prediction Precision and orange bar shows improvement of language model with inverse-folding model, RhoDesign.”High-fitness prediction precision” refers to the fraction of the top 40% predictions that are experimentally determined to confer high RNA fitness, defined as having property within the top 40% of all experimentally screened variants.

We found that both RNA language models and the RNA inverse-folding model can predict mutational effects on RNA function in a zero-shot manner, with performance improving in line with scale laws as data quality, quantity, and model parameters increase. Two recently released RNA language models, AIDO.RNA (1.6B) and RiNALMo (651M), show significant accuracy and robustness compared to other foundation models. Despite having only 37.1M parameters, the inverse-folding model achieved nearly top performance. Notably, all tested DNA language models did not show a significant correlation on the test set.

To verify whether the inverse-folding model could provide positive assistance to the nucleotide language model, we calculated the correlation coefficients between the scores of different models (Fig. 1d). We found that although both the inverse-folding model and the language models exhibited abilities to predict mutational effects on ncRNA function when tested on the RNA DMS dataset, the Spearman correlation coefficient between their scores on the same set of DMS data was very low, which may suggest that the inverse-folding model and the language model capture different dimensions of the relationship between sequence and function. (Extended Data Fig. 2)

Furthermore, we aimed to explore how to integrate these different dimensions of information in a beneficial way to improve the construction of a more comprehensive fitness landscape. Since the principal goal of directed evolution is to identify high-performance variants, we used precision as our primary evaluation metric. We calculated the proportion of sequences in the top 40% scored by both the inverse folding model and the language model where the ground truth score also lies in the top 40%. The results show that compared to using only the language model, incorporating the information captured by the inverse folding model significantly improves the precision score. (Fig. 1f) We also considered additional metrics, such as pLDDT (for structural prediction) and RMSD (for structural similarity), but observed low correlations between these measures and RNA activity across our six datasets. (Extended Data Fig. 3) While there may be more sophisticated ways to combine the inverse folding and language models, even a straightforward multiplication of their normalized scores yields promising results. Building on this, we integrated AIDO.RNA (1.6B) and RhoDesign into a new model, RILLIE, and employed it for RNA directed evolution.

RILLIE, “RNA In Silico Evolution via LLM and Inverse folding”, is a hybrid model combining an RNA language foundation model and an inverse folding model (Fig. 1e). We hypothesize that our model can generate RNAs that conform to evolutionary patterns and natural RNA grammar at the sequence level while maintaining the structural key regions necessary for the original functions. Ideally, it can efficiently evolve any aptamer in just one round without training on specific data. Based on the results from wet-lab experiments, the distribution of generated sequences can be adjusted to exclude harmful mutations and introduce new ones, much like SELEX, making the generated sequence distribution closer to the global optimum in the fitness landscape.

### Directed evolution of Broccoli

To further explore whether our model, RILLIE, which incorporates evolutionary information at the sequence level, can optimize RNA with complex structures more effectively, we focused on designing the fluorogenic Broccoli aptamer, which binds to the small molecule DFHBI-1T, which has been widely used for its fluorescence properties.^31–33^ Structurally, it forms a conserved G-quadruplex motif upon binding to DFHBI-1T, leading to significant fluorescence enhancement, making it a powerful tool for RNA visualization and detection. ^34^However, RNA structure prediction models do not perform well in predicting G-quadruplex motifs.^35^ Therefore, We generated a set of candidate Broccoli sequences and selected 20 sequences for experimental validation to see whether optimization of aptamer systems containing G-quadruplexes could be improved with the help of language models.(Methods)

The number of mutation points is controlled between 3 and 7, as we believe that broccoli has not been well-optimized and therefore has not been trapped in the local optimum. ^34^A moderate amount of mutation can help guide broccoli closer to the global optimal solution. (Fig. 2a) Notably, existing biologically optimized aptamers typically have only 1-2 mutations, as the search space becomes too large with more than 3 mutations. 11 out of 20 aptamers exhibited comparable fluorescence intensity to the wild-type (at least 90% of the wild-type brightness), with 6 of them showing more than a 40% increase in brightness. (Fig. 2b) 13 aptamers that exhibited fluorescent activity were selected for binding affinity testing. All molecules we tested exhibited better affinity than the wild type (Fig. 2b), among which 2 variants (49nt, B15, B16) mutated 6 nucleotides, and 1 variant(B18) mutated 7 nucleotides. B18 with seven mutations still exhibited comparable fluorescence intensity in vitro. After visualizing the structure, most of these mutations are in non-binding regions (Extended Data Fig. 4a). Furthermore, by comparing with previous research on broccoli, it can be observed that the mutation sites selected by our model are mostly located in the variable region.^34^ The B2 variants achieved a 54% increase in fluorescence and a nearly 100% increase in binding affinity. (Fig.2c) These results show that the model has learned structural knowledge to avoid excessive mutations in regions crucial for maintaining structural stability and achieving the multi-property evolution of Broccoli.

**Fig. 2.**
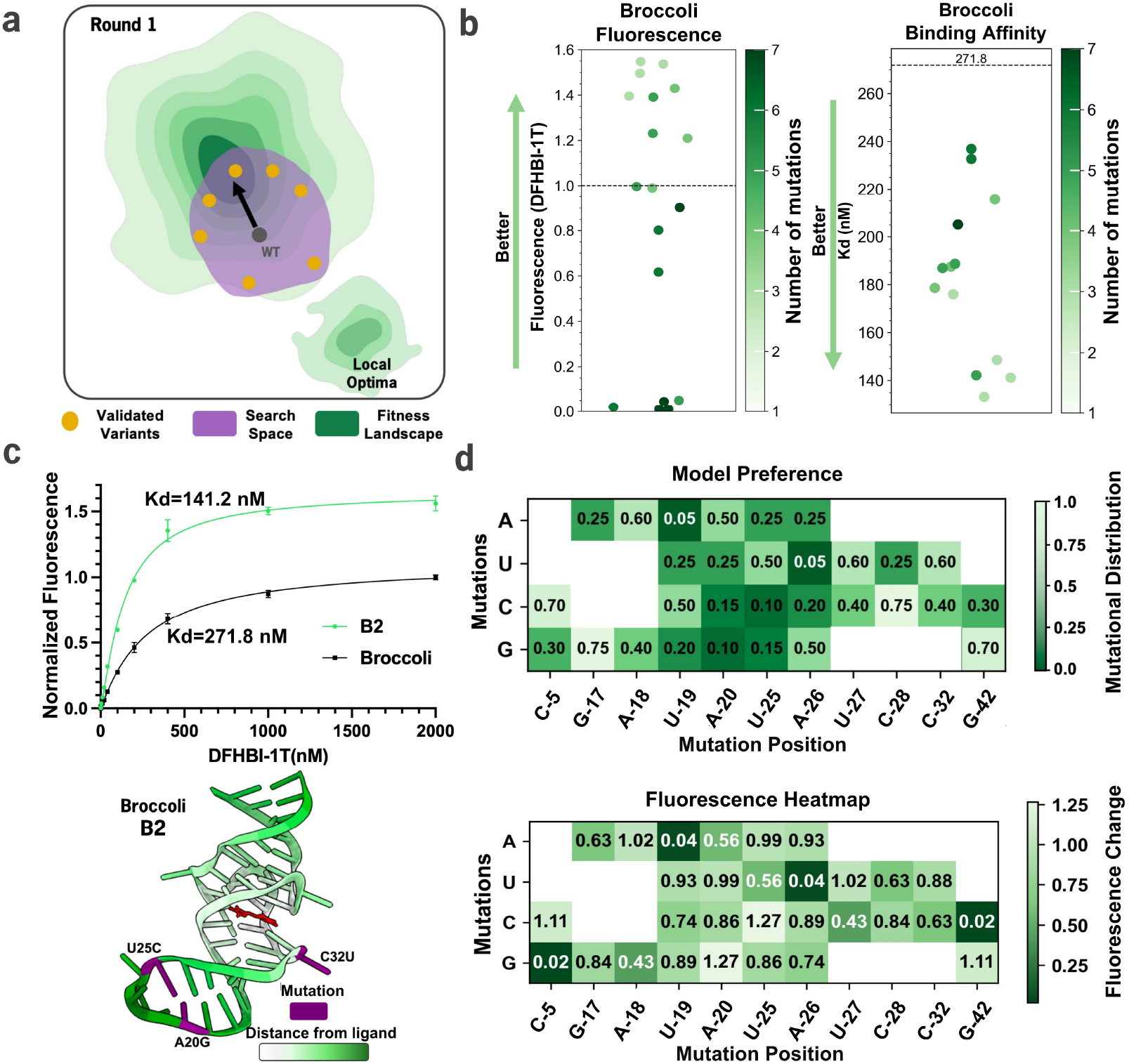
Evolution of fluorescent RNA aptamer with RILLIE improves both affinity and fluorescence. **a**, Schematic diagram of using directed evolution to bring broccoli, which was already close to the local optimum, even closer to the global optimum. **b**, The fluorescence and affinity of the designed variants.13 of 20 designed variants showed fluorescence, among which 9 variants exhibited better fluorescence. All 13 tested variants showed better binding affinity than wildtype. **c**, The EC50 curve of B2 reflects the changes in fluorescence intensity at different dye(DFHBI-1T) concentrations. B2 exhibited a 55% increase in fluorescence intensity in vitro, a 2-fold improvement in binding affinity, and was significantly brighter than the wildtype in vivo. **d**, The relationship between model preference and fluorescence change based on the result of 20 tested variants. The heatmap shows that the model efficiently avoids mutations that cause fluorescence loss in the aptamer (C5G, U19A, A26U, G42C), assigning them very low preference, which aligns with our understanding of the model’s ability in the High-fitness sensitivity analysis (Fig. 1f), where the model is better at selecting mutations that enhance performance and avoiding harmful ones. Additionally, the model is also bold in introducing a variety of mutations at sites that improve fluorescence (U19, A20, U25, A26). Library can be modified based on these information.

To verify whether our model simulates natural selection pressure and preferentially selects mutations beneficial for RNA aptamer directed evolution, we visualized the relationship between the impact of different mutations on fluorescence and the model’s preference for these mutations. We calculated the average fluorescence intensity corresponding to each mutation at every site to determine whether the mutation is beneficial for fluorescence. (Fig.2d) The heatmap shows that the model efficiently avoids mutations that cause fluorescence loss in the aptamer (C5G, U19A, A26U, G42C), assigning them very low preference. This observation supports our high-fitness sensitivity analysis, demonstrating the model’s aptitude for selecting mutations that boost performance while steering clear of detrimental alterations. Additionally, the model is also bold in introducing a variety of mutations at sites that improve fluorescence (U19, A20, U25, A26). These results suggest that the model has indeed learned the principles of natural evolution and is performing directed evolution on the sequence.

To verify whether the natural selection pressure simulated by our model enabled the generated aptamers to sustain the same performance in vivo, we conducted intracellular experiments in human embryonic kidney(HEK) cells with the 13 aptamers that were active in vitro. Firstly, we took images of these 13 sequences in living cells using fluorescence microscopy (Fig.3a) and calculated their fluorescence intensity statistically(Fig.3b). (Methods) For variants with low-point mutations (3-4), they generally showed a significant fluorescence increase, while those with higher-point mutations (5-7) exhibited fluorescence intensities comparable to the wild type. Furthermore, we used FACS to perform more precise statistical analysis and measurement of the fluorescence intensity in the cells. Fluorescence-Activated Cell Sorting (FACS), a bench mark flow cytometry-based technique, enables precise single-cell quantification of fluorescence intensity in living cell populations. The in vivo performance of all variants was also consistent with their extracellular results. (Fig.3c) These results prove that the simulated natural selection pressure indeed evolved the generated aptamers to fold more efficiently, thus enhancing their cellular functionality instead of selecting only for binding affinity like SELEX.

**Fig. 3.**
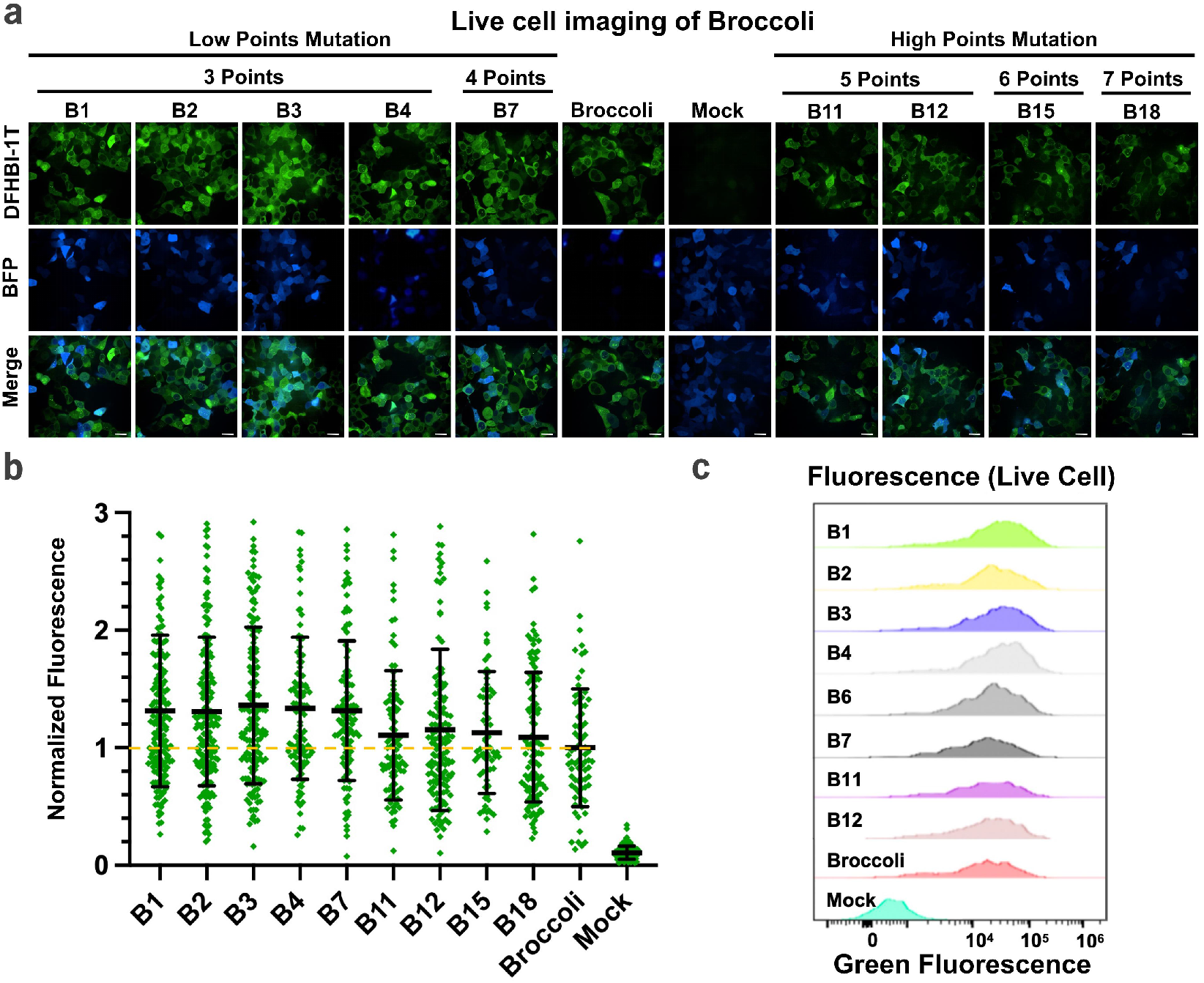
Evolution of RNA aptamer with RILLIE improves fluorescence in HEK293T cells. **a**, Confocal imaging of live HEK293T cells expressing the selected low-mutation (B1, B2, B3, B4, B7) and high-mutation (B11, B15, B18) variants. HEK293T cells transfected with plasmid expressing wild-type Broccoli or the mock plasmid were used as the controls. The cells were cotransfected with a plasmid expressing blue fluorescent protein to distinguish transfected cells from nontransfected ones. Scale bars, 20 μm. **b**, Quantitative analysis of the fluorescence of Broccoli-DFHBI in individual HEK293T cells in **(a)**. The data represent the mean ± s.d (n=50 cells). **c**, Flow cytometry histograms of the fluorescence distributions for different Broccoli variants. The intracellular environment is more complex, and fluorescent RNA is often under expressed, which imposes stricter requirements on affinity.Eight variants exhibit higher fluorescence than wildtype.

### Pepper Round 1

To further validate the generalizability of our model, we attempted to optimize another fluorescent aptamer. As a starting point, we considered the pepper variant with the highest fluorescence and binding affinity, which has been artificially optimized. Fluorogenic Pepper aptamer^29^, a small molecule(49nt), has enormous practical value and is continually being researched and modified by scientists^36 37,38^but haven’t been structurally resolved (the pepper we designed).

In the first round of evolution, the number of mutation points is controlled between 4 and 6 to explore the fitness landscape around pepper with low-point mutations (Fig.4a). (Methods) Upon testing the synthesized aptamers for fluorescence in the presence of HBC-530, we found that 15 of the 20 optimized aptamers exhibited higher fluorescence intensity than the wildtype, with 9 of them showing an increase of more than 30% (Fig. 4b). Additionally, we tested the fluorescence in the presence of another dye, HBC-620, and 14 of the 20 optimized aptamers exhibited higher fluorescence intensity. (Extended Data Fig. 8).

**Fig. 4.**
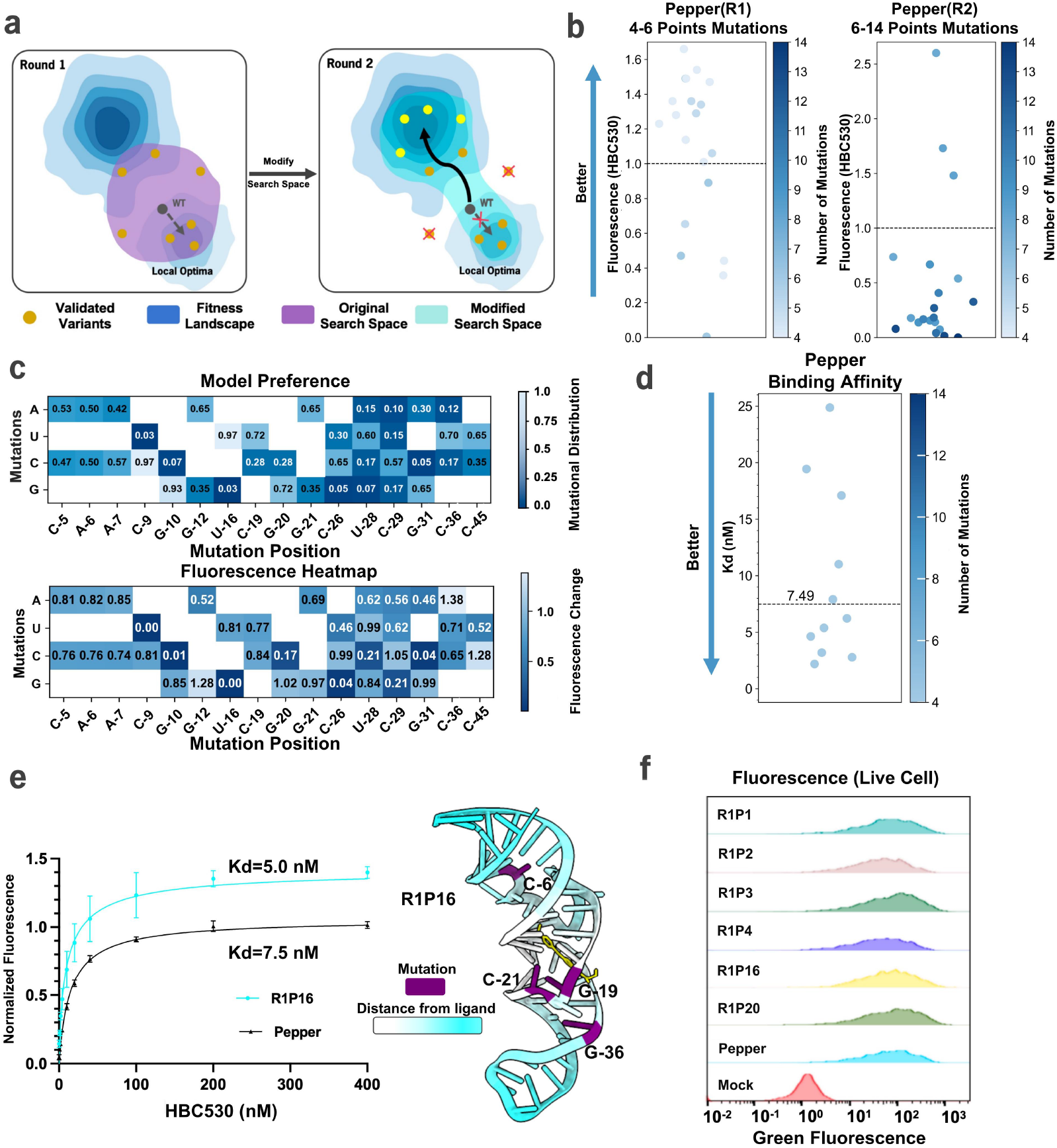
Evolution of fluorescent RNA aptamer with RILLIE improves sequence novelty, affinity and fluorescence in HEK293T cell for different dyes. **a**, Schematic diagram of the strategy to modify the search space for better variants. The yellow dots represent variants tested in wet-lab experiments. Since nucleotide foundation models excel at identifying high-fitness mutations, we can discard harmful mutations and introduce new mutations based on test results. Discarding harmful mutations can increase the success rate, while introducing new mutations can help direct evolution escape local optima and discover global optima. This approach enables efficient directed evolution without retraining the model. **b**, Fluorescence improvements across two rounds for HBC620 and HBC530. Strip plots show variant fluorescence changes vs. wild-type Pepper. In round 1 (50 variants, 3–6 mutation points each), 24 variants surpassed 30% improvement for at least one dye. In round 2 (20 variants, 6–14 mutations), HBC530 fluorescence increased by up to 2.6-fold, and one 12-mutation variant remained strong for HBC620. **c**, Relationship between model preference and fluorescence change based on the result of 40 tested variants. The heatmap shows that the model efficiently avoids mutations that cause fluorescence loss in the aptamer (C_9_U, G_10_C, U_16_G, G_20_C, C_26_G, G_31_C), assigning them very low preference. This aligns with our understanding of the model’s ability in the High-fitness sensitivity analysis (Fig. 1f), where the model is better at selecting mutations that enhance performance and avoiding harmful ones. Library can be modified based on this information. **d**, Trade-offs between affinity and fluorescence. Some variants match or exceed Pepper in terms of the binding affinity to HBC530. However, more extensive mutations often degrade affinity despite boosting fluorescence. **e**, The EC_50_ curve of R1P16 illustrates fluorescence intensity at different dye concentrations. Compared with wild-type Pepper, R1P16, which contains four nucleotide mutations with two located inside the binding pocket and two located outside the binding pocket, exhibited a 1.5-fold increase in the binding affinity to HBC530, and maintained higher fluorescence intensity across all dye concentrations. **f**, Flow cytometry histograms of the fluorescence distributions for different Pepper variants. Five variants exhibit higher fluorescence than wildtype.

To test whether our method can further optimize aptamers with nanomolar-level affinity and improve their binding affinity, we tested the affinity of aptamers with enhanced fluorescence using HBC-530 and HBC-620. We found that 7 of these molecules exhibited improved affinity, while 11 showed comparable affinity with HBC-530 (with a Kd change within a 5-fold range).(Fig.4b) Notably, 5 of the generated variants showed an almost 3-fold increase in binding affinity with HBC-620.( Supplementary Data 2) These results indicate the successful directed evolution of pepper for both dyes.

### Pepper Round 2

We hypothesize that since the Pepper aptamer has already undergone extensive optimization by previous researchers,^39–41^ any remaining potential for improvement may be constrained by a local optimum^38,42^—a common challenge in wet lab based directed evolution. We are interested in whether performing a second round of zero-shot evolution using RILLIE will allow us to escape potential local maxima traps and discover an aptamer with better performance. (Fig.4a) Once again, we initiated the directed evolution process using the wild-type sequence as our starting point. This time, we increased the number of mutations from 4–6 to 6–14. Using the same pipeline, we generated a set of candidate Pepper sequences and selected 20 sequences for experimental validation. (Methods)

5 out of the 20 aptamers exhibited higher fluorescence intensity (using the same criterion as the first round). One variant (6 points mutation) showed a 2.6-fold increase in fluorescence intensity when binding with HBC-530. (Fig.4b) Another aptamer had 13 mutations but still maintained the same fluorescence intensity as the wildtype with HBC-620. (Extended Data Fig.8)

We calculated the average fluorescence intensity corresponding to each mutation at every site to determine whether the mutation is beneficial for fluorescence with two rounds of fluorescence data. (Fig.4c) The heatmap shows that the model efficiently avoids mutations that cause fluorescence loss in the aptamer (C9U, G10C, U16G, G20C, C26G, G31C), assigning them very low preference. These results (pepper and broccoli) suggest that the model has indeed learned the principles of natural evolution and is performing directed evolution on the sequence.

Intracellular experiments were conducted on variants with improved fluorescence in vitro. FACS was employed to achieve precise single-cell fluorescence quantification while ensuring that only live cells were analyzed. We found four variants exhibit better *in vivo* fluorescence than wild type. (Fig.4f) It is worth mentioning that the full length of Pepper is 49nt, which means our sequence similarity is as low as 0.7, comparable to the sequence similarity in some de novo design settings^43^.

## Discussion

In this study, we have demonstrated the power of language models augmented with inverse-folding models to efficiently guide the directed evolution of RNA aptamers in a zero-shot fashion, achieving significant improvements in fluorescence *in vivo* and binding affinity. By simulating natural selection pressure, our approach bridges the gap between in vitro performance and in vivo functionality, producing aptamers that exhibit robust folding and cellular performance reminiscent of naturally evolved counterparts. Furthermore, our method was able to escape potential local maxima traps and discover variants with better performance by conducting a second round of evolution, Importantly, this method bypasses the need for task-specific training data, providing a more scalable and less resource-intensive approach to RNA optimization.

One of the key strengths of our approach lies in the integration of language models and inverse-folding models. Compared to using language models alone for RNA directed evolution^44^, our model has the advantage of capturing structural information, which may help maintain the structural integrity of crucial regions. In contrast to using inverse-folding models alone for RNA directed evolution^8^, our model has been trained on a large corpus of sequences, allowing it to learn the patterns of natural evolution. By simulating natural selection pressure, the designed sequences are more likely to align with natural evolutionary patterns at the sequence level, thus enhancing the likelihood that the optimized RNA will function effectively in vivo. In the case of Broccoli, our method avoids excessive mutations that could destabilize essential RNA structures, with mutations predominantly occurring in variable regions. In the case of Pepper, not biased by traditional structural hypotheses, many beneficial mutations were structurally remote from the ligand-binding domain, which are typically excluded from traditional directed evolution strategies. These mutations were able to enhance fluorescence and binding affinity without compromising the RNA’s structural stability.

Our method is theoretically applicable to any RNA, though we have only validated the model’s capabilities on two RNA aptamer systems. In our benchmarking and high fitness prediction precision assessments, the model performed comparably well on tRNA and ribozyme datasets, demonstrating the robustness of the approach across different RNA types.

We anticipate that, as these models continue to improve, the use of nucleotide foundation models for zero-shot RNA optimization design will become a key part of RNA engineering, revolutionizing the way we approach RNA-based applications. In our model’s benchmark, we validated the scaling law and found that larger models trained on higher-quality datasets tend to show better generalization performance. This observation suggests that increasing the quantitiy and quality of the training data and the scale of models could further enhance the model’s capability to predict RNA structure and mutational effects and evolve diverse RNAs across various functions.

## Methods

### Model description and zero-shot fitness prediction of sequences

In this work, we propose RILLIE – “RNA In Silico Evolution via LLM and Inverse folding”, a hybrid model combining an RNA language foundation model and an inverse folding model for zero-shot fitness prediction of RNAs undergoing directed evolution. This approach eliminates the need for explicit fitness data during model training. By combining the evolutionary sequence patterns captured by the large language model and the structural compatibility predicted by the inverse folding model, we create a composite fitness score to identify promising candidate sequences for experimental validation. We adopt AIDO.RNA, the RNA language model with the largest number of parameters, as the language foundation model to capture natural evolutionary information and assess sequence plausibility. Simultaneously, we select RhoDesign, the current state-of-the-art and most stable RNA inverse folding model, as the structure foundation model to evaluate the possibility of RNA sequences with given structures.

The language foundation model (AIDO.RNA) aims to evaluate the plausibility of the evolved sequence ***s*’** under natural evolutionary patterns. AIDO.RNA is trained on a large-scale RNA sequence dataset to capture potential evolutionary patterns of RNA sequences. The input to the language model is the RNA sequence 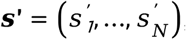, where 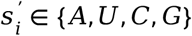 denotes the nucleotide type at the *i*-th position and represents the number of nucleotides in the RNA. AIDO.RNA computes the probability of ***s* ‘** auto-regressively:

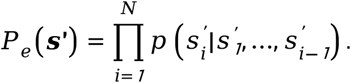

Here, 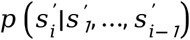 represents the probability of the *i*-th nucleotide being 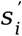 given the preceding *i*− *1*bases. *P*_*e*_ (***s*’)** quantifies the global evolutionary plausibility of the sequence under the language model. A higher *P*_*e*_ (***s*’)** indicates that the sequence ***s*’** aligns more closely with the natural RNA evolutionary patterns learned by AIDO.RNA.

The inputs to the structure foundation model include the RNA tertiary structure ***C*** ∈ ℝ ^*N*×*7*×*3*^ and secondary structure information (nucleotide contact map) extracted from the tertiary structure. Each nucleotide is represented by the 3D coordinates of its seven backbone atoms: *C 4*’, *C 1*’, *N1, C 2, C 5*’, *O5*’ and *P*. The RNA secondary structure is extracted from the tertiary structure using DSSR42 default settings and encoded as a feature vector. This secondary structure feature is combined with the tertiary structural coordinates as input to RhoDesign to enhance the model’s structural representation capability. The structural foundation model learns the probability distribution of RNA sequences ***S***= (*S*_*1*_, …,*S*_*N*_) conditioned on the structure ***C*** through an autoregressive approach, formalized as:

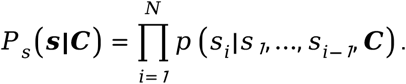

For a directed-evolved sequence 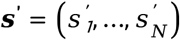 and its target structure ***C*’**, we can compute:

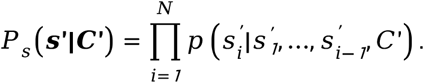

This probability *P*_*s*_(***s*’**| ***C*’**) represents the likelihood of sequence ***s*’** occurring under the condition of structure ***C*’**. We assume that the wild-type RNA structure already satisfies functional requirements; thus, serves as an indicator of the compatibility between the evolved sequence and the target structure, indirectly reflecting its potential structural fitness.

To systematically evaluate the fitness of RNA sequences generated through directed evolution, we define an integrated scoring function:

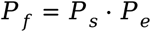

Here, *P*_*s*_(***s*’**| ***C*’**)is derived from the inverse-folding model, originally output as a log-likelihood reflecting the probability of sequence ***s*’** occurring given a fixed structure ***C*’**. We assume the target structure ***C*’** corresponds to the wild-type RNA conformation, which satisfies functional requirements; thus,*P*_*s*_ serves as an indicator of the evolved sequence’s compatibility with this structure, indirectly assessing its structural fitness. Similarly, *P*_*e*_ (***s*’)** is obtained from the language model, also initially computed as a log-likelihood, representing the prior probability of sequence ***s*’** in a natural evolutionary context. In our implementation, both log-likelihoods are exponentiated and normalized to the [0, 1] range, ensuring they are comparable probabilities suitable for multiplication. This scoring approach is inspired by the “Product of Experts” (PoE) framework by Geoffrey E Hinton^45^, where the product of normalized probabilities from independent models yields a joint distribution capturing both structural compatibility and evolutionary plausibility.

The combined score *P*_*f*_ = *P*_*s*_⋅*P*_*e*_ integrates these two dimensions into a single metric, prioritizing sequences that align with the functionally validated wild-type structure while retaining natural evolvability. This practice of multiplying scores is reasonable and has been similarly employed in protein design, such as in Plug and Play Directed Evolution (PPDE)^46^. By selecting sequences with the highest *P*_*f*_, we aim to identify RNA variants with both good structural fidelity and evolutionary likelihood, enhancing their potential for functional validation in downstream experiments.

Ultimately, RNA sequences ranked by *P*_*f*_ will be subjected to wet-lab experiments to confirm their functional performance in a biological setting. This strategy leverages the strengths of pre-trained models, integrating structural and evolutionary information through a normalized probability product, and provides a theoretically grounded and experimentally actionable framework for RNA sequence optimization.

### Generation of Broccoli sequences

We employed RILLIE to optimize the Broccoli system through a two-phase strategy: single-site mutation screening and combination optimization. In the first phase, all nucleotide positions were systematically evaluated for three potential mutation types using this dual-model computational framework. The RhoDesign model quantified evolutionary preference, while AIDO.RNA predicted structural plausibility. Mutations were prioritized based on their ability to outperform the wild type in both metrics. At each site, if one mutation type improved both scores, all dual-improvement variants were retained as candidates; otherwise, mutations enhancing either metric were included. To minimize redundancy, each site was calculated the composite score (using the formula detailed in the previous section) reflecting the highest-performing mutation variant. Sites were ranked by descending composite scores, with the top 11 selected for subsequent combination optimization.

During the second phase, mutations from candidate sites were systematically combined to generate mutant libraries spanning 3–7 modified sites. Each resulting sequence was again evaluated by the composite score, integrating evolutionary preference and structural plausibility. The 20 top-ranked sequences were then chosen for experimental validation. This hierarchical approach enabled thorough assessment of single-site changes and systematic exploration of potential synergistic interactions, ultimately identifying Broccoli variants with enhanced functional performance and structural integrity.

### Generation of Pepper Sequences

The evolutionary workflow for the Pepper system followed the same two-phase strategy as the Broccoli system—single-site mutation screening followed by combination optimization—but incorporated iterative modifications to address its unique optimization challenges. Given Pepper’s well-characterized properties from prior studies, we hypothesized that restricting mutations to fewer sites might trap the system in local optima, limiting performance gains. To test this, we implemented a two-round evolutionary strategy: the first round introduced 4–6 combination mutations, while the second round expanded exploration to 6–14 sites, excluding combinations that abolished activity in the initial phase and introducing novel mutation sites. All variants were assessed using the composite score integrating evolutionary preference and structural plausibility, with the top 20 sequences selected for wet-lab validation. This iterative approach balanced the preservation of functional RNA structures with aggressive exploration of high-dimensional mutational space, explicitly investigating whether increased site diversity could overcome local optima and enhance performance, ultimately refining the evolution of the Pepper system.

### Curating diverse ncRNA scanning mutagenesis data

Given that no well-established benchmark datasets exist for ncRNA function prediction, we curated the literature for examples of ncRNA mutational scanning experiments. We obtained the following datasets: a ribozyme DMS by Andreasson et al.^30^,a tRNA DMS by Domingo et al.^47^,a tRNA DMS by Guy et al.^28^, a fluorescent RNA aptamer DMS by Chen et al.^29^, a fluorescent RNA aptamer DMS by Zuo et al.^10^, a fluorescent RNA aptamer DMS by Jiang et al.^11^. The dataset requires that the internal sequence lengths be consistent, as the sequence length input for the inverse-folding model must match the given three-dimensional structure. For the ribozyme DMS dataset by Andreasson et al. (25), we removed sequences with lengths differing from the original sequence and normalized the fitness scores to the range of 0 to 1. We selected data with fitness greater than 0.06. Some sequences correspond to multiple fitness values under different experimental conditions. To bridge the gap between in vivo and in vitro data distributions, we selected the data most representative of in vivo performance. During the dataset search, we also found some ncRNA DMS datasets with varying sequence lengths, which might be useful for future research. These include a ribozyme dataset (1w+) [https://datadryad.org/stash/dataset/doi:10.5061/dryad.nm1189t] and a sRNA dataset (2000+) [https://datadryad.org/stash/dataset/doi:10.5061/dryad.3rq8q88]. Detailed information about the datasets is provided in **Supplementary Information**.

### Benchmarking diverse nucleotide foundation models with scanning mutagenesis data

We used DMS datasets to benchmark nucleotide foundation models based on their ability to predict mutational effects on ncRNA function. Nucleotide foundation models consist of three types of models: RNA language model, genomic language model, and inverse-folding model, which are proposed after 2022 that performed well on previous benchmark tests. These models included AIDO.RNA^22^, RiNALMo^23^, RNAFM^20^, RNAMSM^24^(RNA language models), Evo^17^, Nucleotide Transformer^25^, GENA-LM^26^ and GROVER^27^(genomic language models)and the RhoDesign^8^(RNA inverse-folding model).

The **RNA language model** and **DNA language model** are both trained on large datasets of nucleotide sequences using a masking learning approach. These models learn the statistical dependencies and patterns within RNA and DNA sequences by predicting masked nucleotides based on their context. This training process allows the models to capture valuable information about the natural evolutionary strategies that shape these sequences.

Natural evolution explores a vast landscape of possible sequences, searching for rare mutations that are desirable. By learning from this evolutionary process, these models gain insight into how sequences evolve and adapt over time. This knowledge of natural evolutionary strategies can then inform artificial evolution, guiding the design and optimization of sequences for specific biological functions or properties.

The **inverse-folding model** is a computational approach used to predict RNA or protein sequences that will fold into a specific desired three-dimensional structure. Unlike traditional folding models, which predict the structure given a sequence, inverse-folding models work in reverse, generating potential sequences that can achieve a specified target structure. This is particularly useful for designing novel biomolecules with desired functionalities or for engineering RNA/protein molecules with specific folding patterns.

To benchmark our inverse-folding model, RhoDesign, which was previously a structure-to-sequence model, we modified its input and output. The input consists of the 3D structure of the wildtype of each system, the secondary structure, and the sequence for which the fitness needs to be predicted. The output is the likelihood of this sequence predicted by the model. We calculated the Spearman correlation coefficient between the likelihood and the fitness of the true sequence. The 3D structure was predicted by AlphaFold3, and the secondary structure was predicted by RhoFold.

To benchmark the RNA language model and the genomic language model, we selected the most up-to-date model weights as of the time we conducted the experiment, and computed the Spearman correlation between the experimental fitness values and the sequence pseudolikelihood (for masked language models).

### Benchmarking diverse metrics with scanning mutagenesis data

**plDDT (per-residue plDDT)** is a metric used in protein structure prediction. It represents the per-residue confidence score of a predicted protein structure, with values ranging from 0 to 100. Studies have shown that proteins with higher plDDT scores tend to have better functionality. With the open-source release of AlphaFold3 in 2024, we used AF3 to perform structural predictions on five datasets across six benchmarks, predicting over 4,000 structures. This effort also provides a foundational dataset for future RNA research from a structural perspective. We calculated the Spearman correlation coefficient between the plDDT score of each sequence in the datasets and their experimental fitness. However, we found that the correlation coefficients were generally low, indicating that plDDT is not a good metric for reflecting experimental fitness. Therefore, to conserve resources, we did not compute the 3D structure for the ribozyme DMS dataset by Andreasson et al. (2025).

**RMSD (Root Mean Square Deviation)** is a widely used measure in structural biology to quantify the average distance between corresponding atoms, typically backbone atoms, of two molecular structures.

To investigate whether the inverse-folding model captures the mapping between sequence and experimental fitness simply by calculating the structural similarity (RMSD) between variants and the wildtype sequence, we performed structural predictions for five benchmark datasets using RhoFold and AlphaFold3. We selected the predicted structures with the highest confidence and compared them with the corresponding wildtype structures (predicted using AF3) to calculate RMSD. Subsequently, we computed the correlation coefficient between RMSD and experimental fitness. The results showed a very low correlation, indicating that the inverse-folding model does not rely solely on structural similarity to score sequences.

Detailed information about the benchmark data is provided in **Supplementary Data 1**.

### Visualize relationship between mutations and fluorescence

Unlike traditional RNA directed evolution, which involves single-point mutations, RILLIE guided evolution involves 4-14 mutation sites. It is difficult to determine the effect of a specific mutation on experimental fitness from the relationship between the combination of these mutation sites and experimental fitness, due to synergistic effects and epistasis. We would like to use a method to visually represent this.

We calculated the average fluorescence intensity corresponding to different nucleotide mutations at each position of a sequence, using this as a numerical assessment of whether a mutation is beneficial or detrimental to fluorescence. We then plotted a heatmap to visually reflect this. For example, if the first nucleotide of a wildtype sequence is A, and in the variants tested experimentally, it is mutated to U and C, there are three possible variations for the first nucleotide: A-1-A, A-1-U, and A-1-C. The average fluorescence intensity for each mutation is calculated by dividing the fluorescence intensity of the corresponding sequence by the number of corresponding sequences. This method is particularly useful for identifying harmful mutations, as these mutations will cause a decrease in fluorescence intensity regardless of the mutations at other positions. As a result, the average fluorescence intensity for these mutations will be close to 0 in the final plot.

## Acknowledgements

Y.Y. was supported by the National Key Research and Development Program of China (2022YFC3400100), NSFC (32121005, 92357308, 22437001). S.Z. was supported by supported by the National Natural Science Foundation of China [62041209], Natural Science Foundation of Shanghai [24ZR1440600], the Young Elite Scientists Sponsorship Program by CAST [2023QNRC001], the Science and Technology Commission of Shanghai Municipality [24510714300]. S. Z. acknowledges funding from the Asian Young Scientist Fellowship.

## Author contributions

S.Z. conceived and supervised research. Y.H. conceived research, developed RILLIE, performed or directed all experiments, and wrote the paper. J.X.C.and R.M. and Z.Z. performed computational experiments. B.Z. performed experiments and analyses. X.C. and Y.Y. were involved in the discussion and contributed to the organization. All authors assisted with manuscript editing.

## Data availability

Detailed results of our benchmarks are in Supplementary Data 1.

Generated and tested sequences of Broccoli and Pepper, in addition to Kd, fluorescence activity are in Supplementary Data 2.

FACS data of Broccoli and Pepper are in Supplementary Data 3.

The fluorescence data in living cells are in Supplementary Data 4.

The information of primers are in Supplementary Data 5.

## Code availability

RILLIE is available at https://github.com/GENTEL-lab/RILLIE.

**Extended Data Fig. 1.**
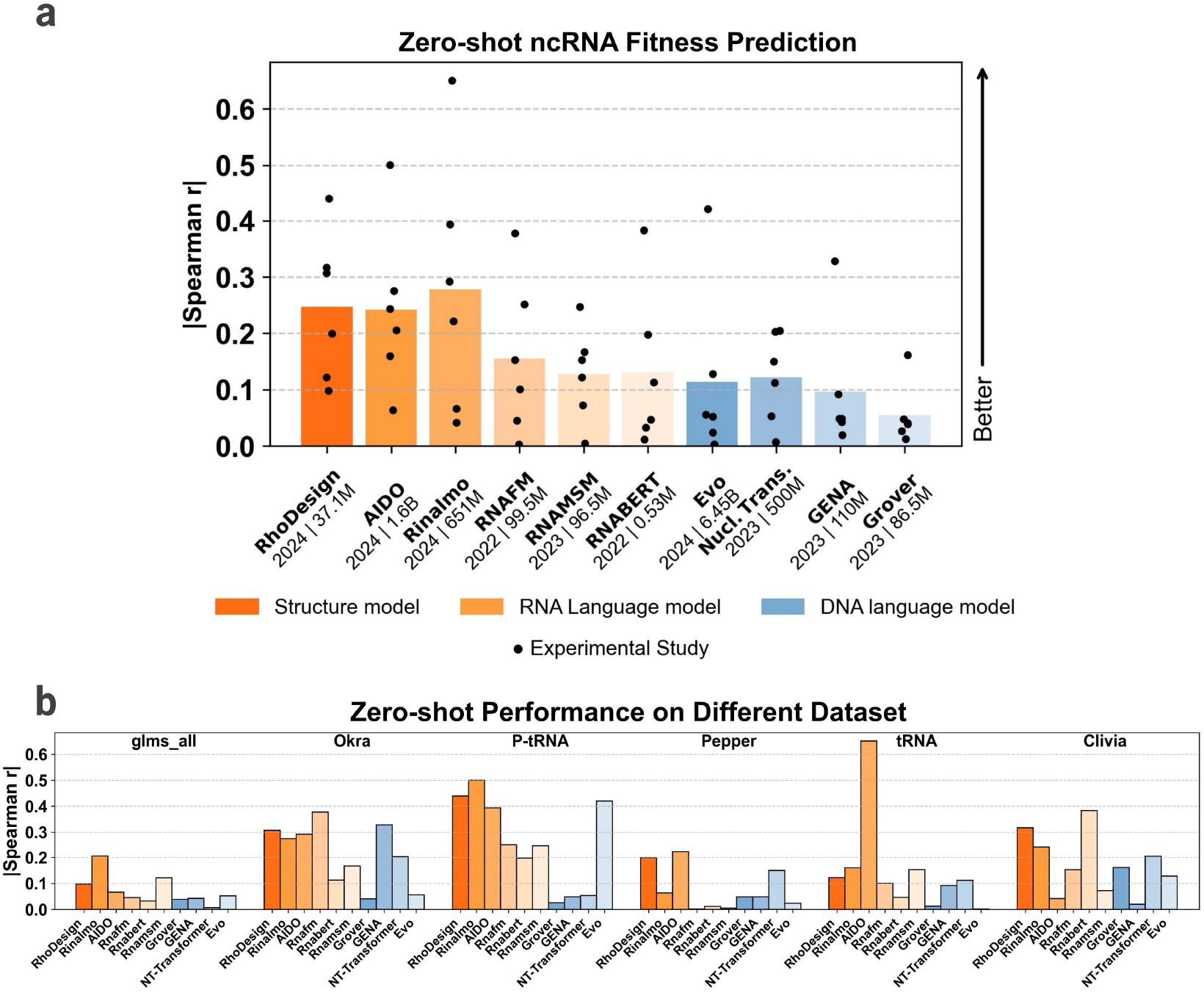
The unsupervised log-likelihood of the nucleotide foundation model exhibits certain correlation with RNA fitness. a. The absolute Spearman correlation coefficient between the inverse-folding model, RNA language model, and DNA language model and fitness (DMS info shown in Table 1). Bar height indicates the mean; each dot indicates a different DMS study. Nucl.-Trans., Nucleotide Transformer. b. The specific absolute correlation coefficients of various nucleotide foundation models across different datasets. (Detailed data in **Supplementary Data 1**).

**Extended Data Fig. 2.**
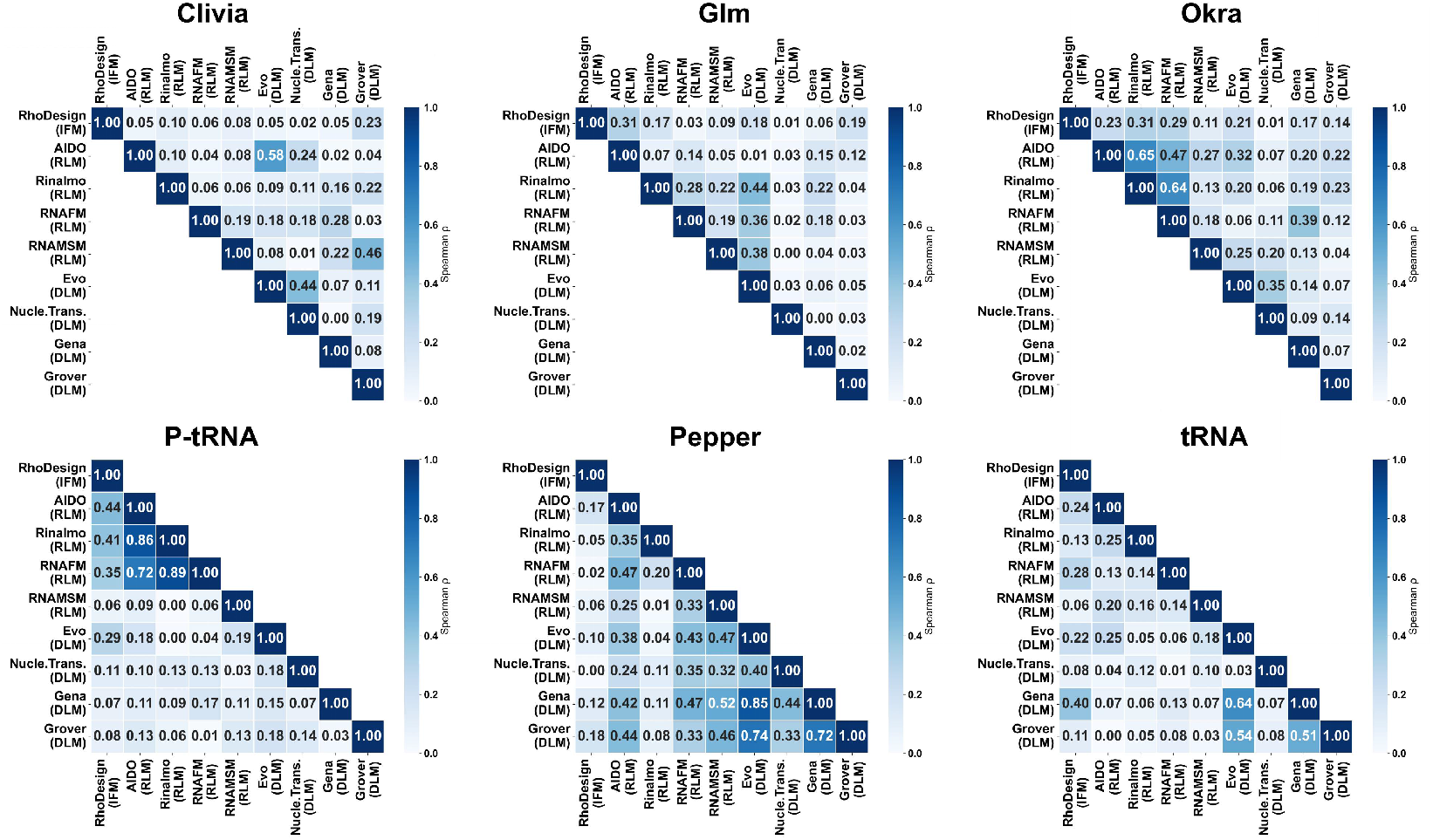
The Spearman Correlation between different models across six datasets. The correlation between RNA models and RNA models, as well as the correlation between DNA models and DNA models, is higher than the correlation between RNA models and DNA models. This is rational because models of the same type are more likely to reach a consensus. The inverse-folding model shows generally low correlation with most models (maximum 0.44). However, after cross-scoring, there is a significant improvement in High-fitness Prediction Precision, suggesting that the inverse-folding model and language models capture evolutionary information from sequence and structure dimensions, respectively, enabling more efficient directed RNA evolution.Notably, for our optimized Pepper, the inverse-folding model exhibits very low correlation with most models (maximum 0.18). Nevertheless, after cross-scoring with RNA language models, there is a significant improvement in the High-fitness Prediction Precision of the language model. (Detailed data in **Supplementary Data 1**).

**Extended Data Fig. 3.**
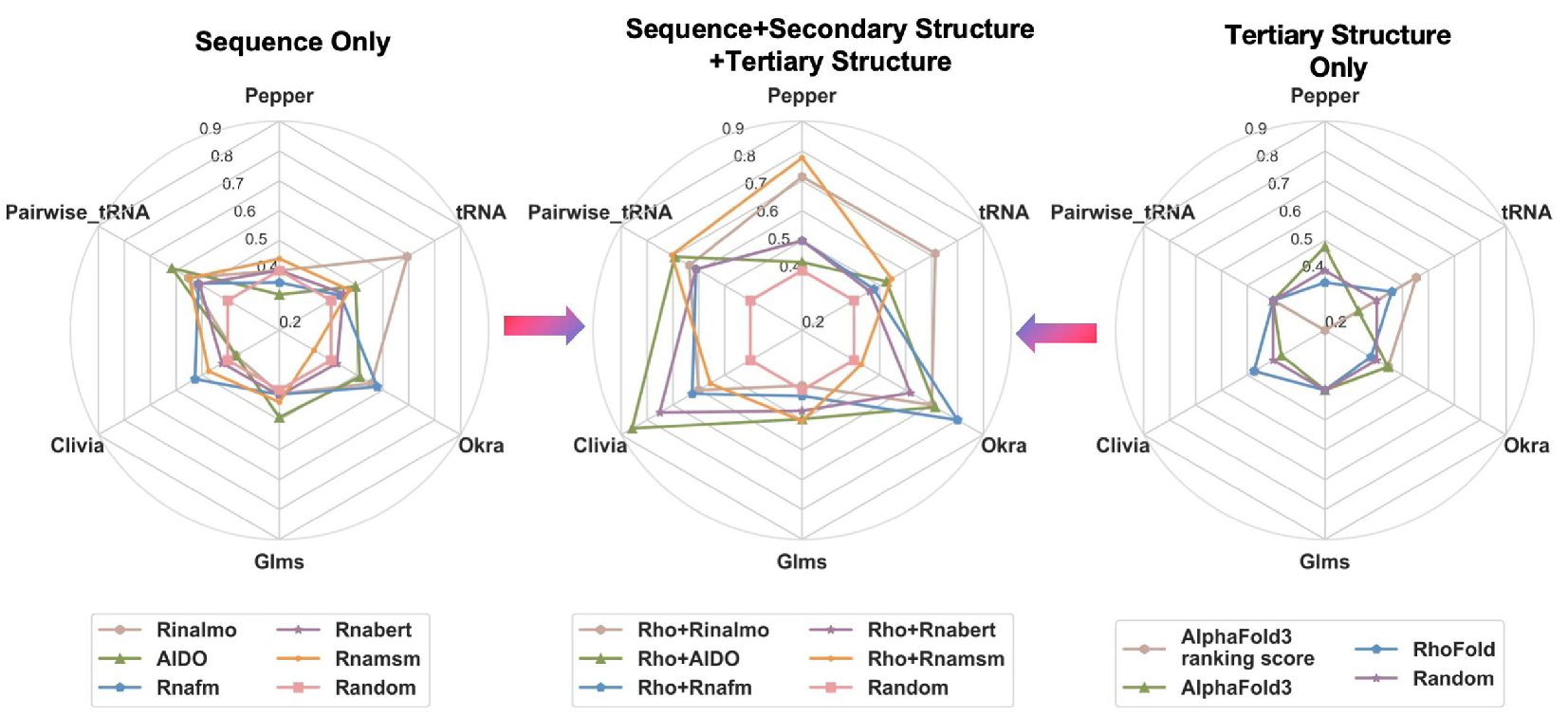
Multimodal input improves language model performance. High-fitness prediction precision of the inverse-folding model in combination with different RNA LLMs(IF+LLM), purely sequence-based(LLM), and purely structure-based (plDDt) approaches.High-fitness sensitivity analysis reveals that multimodal input improves language model performance compared with sequence-only input across 6 RNA from diverse RNA families. “High-fitness prediction precision” refers to the fraction of the top 40% predictions that are experimentally determined to confer high protein fitness, defined as having property within the top 40% of all experimentally screened variants. The left picture examines the relationships between scores solely using the language model and fluorescence, the middle picture represents the relationship of the intersection of RhoDesign scores (reflecting 2D and 3D structural similarity) and language model scores (reflecting sequence evolutionary information) and fluorescence, and the right picture examines the relationships between structural similarity (RMSD), AlphaFold3-derived plDDT and fluorescence. (Detailed data in **Supplementary Data 1**).

## Reference

1. Spitale, R. C. & Incarnato, D. Probing the dynamic RNA structurome and its functions. Nat. Rev. Genet. 24, 178–196 (2023).

2. Wang, X.-W., Liu, C.-X., Chen, L.-L. & Zhang, Q. C. RNA structure probing uncovers RNA structure-dependent biological functions. Nat. Chem. Biol. 17, 755–766 (2021).

3. Ganser, L. R., Kelly, M. L., Herschlag, D. & Al-Hashimi, H. M. The roles of structural dynamics in the cellular functions of RNAs. Nat. Rev. Mol. Cell Biol. 20, 474–489 (2019).

4. Jasinski, D., Haque, F., Binzel, D. W. & Guo, P. Advancement of the Emerging Field of RNA Nanotechnology. ACS Nano 11, 1142–1164 (2017).

5. Pardi, N., Hogan, M. J., Porter, F. W. & Weissman, D. mRNA vaccines — a new era in vaccinology. Nat. Rev. Drug Discov. 17, 261–279 (2018).

6. Townshend, B., Kennedy, A. B., Xiang, J. S. & Smolke, C. D. High-throughput cellular RNA device engineering. Nat. Methods 12, 989–994 (2015).

7. Su-Tobon, Q. et al. CRISPR-Hybrid: A CRISPR-Mediated Intracellular Directed Evolution Platform for RNA Aptamers. Nat. Commun. 16, 595 (2025).

8. Wong, F. et al. Deep generative design of RNA aptamers using structural predictions. Nat. Comput. Sci. (2024) doi:10.1038/s43588-024-00720-6.

9. Hou, Q. & Jaffrey, S. R. Synthetic biology tools to promote the folding and function of RNA aptamers in mammalian cells. RNA Biol. 20, 198–206 (2023).

10. Zuo, F. et al. Imaging the dynamics of messenger RNA with a bright and stable green fluorescent RNA. Nat. Chem. Biol. 20, 1272–1281 (2024).

11. Jiang, L. et al. Large Stokes shift fluorescent RNAs for dual-emission fluorescence and bioluminescence imaging in live cells. Nat. Methods 20, 1563–1572 (2023).

12. Keefe, A. D., Pai, S. & Ellington, A. Aptamers as therapeutics. Nat. Rev. Drug Discov. 9, 537–550 (2010).

13. DeRosa, M. C. et al. In vitro selection of aptamers and their applications. Nat. Rev. Methods Primer 3, 54 (2023).

14. Song, L., Segal, E. & Xing, E. Toward AI-Driven Digital Organism: A System of Multiscale Foundation Models for Predicting, Simulating and Programming Biology at All Levels.

15. Hayes, T. et al. Simulating 500 million years of evolution with a language model. Preprint at 10.1101/2024.07.01.600583 (2024).

16. Shanker, V. R., Bruun, T. U. J., Hie, B. L. & Kim, P. S. Unsupervised evolution of protein and antibody complexes with a structure-informed language model. Science 385, 46–53 (2024).

17. Nguyen, E. et al. Sequence modeling and design from molecular to genome scale with Evo. Science 386, eado9336 (2024).

18. Hie, B. L. et al. Efficient evolution of human antibodies from general protein language models. Nat. Biotechnol. 42, 275–283 (2024).

19. Wang, N. et al. Multi-purpose RNA language modelling with motif-aware pretraining and type-guided fine-tuning. Nat. Mach. Intell. 6, 548–557 (2024).

20. Chen, J. et al. Interpretable RNA Foundation Model from Unannotated Data for Highly Accurate RNA Structure and Function Predictions. Preprint at http://arxiv.org/abs/2204.00300 (2022).

21. Schneider, B. et al. When will RNA get its AlphaFold moment? Nucleic Acids Res. 51, 9522–9532 (2023).

22. Zou, S. et al. A Large-Scale Foundation Model for RNA Function and Structure Prediction. Preprint at 10.1101/2024.11.28.625345 (2024).

23. Penić, R. J., Vlašić, T., Huber, R. G., Wan, Y. & šikić, M. RiNALMo: General-Purpose RNA Language Models Can Generalize Well on Structure Prediction Tasks. Preprint at http://arxiv.org/abs/2403.00043 (2024).

24. Zhang, Y. et al. Multiple sequence-alignment-based RNA language model and its application to structural inference.

25. Dalla-Torre, H. et al. Nucleotide Transformer: building and evaluating robust foundation models for human genomics. Nat. Methods (2024) doi:10.1038/s41592-024-02523-z.

26. Fishman, V. et al. GENA-LM: A Family of Open-Source Foundational Models for Long DNA Sequences.

27. Sanabria, M., Hirsch, J., Joubert, P. M. & Poetsch, A. R. DNA language model GROVER learns sequence context in the human genome. Nat. Mach. Intell. 6, 911–923 (2024).

28. Guy, M. P. et al. Identification of the determinants of tRNA function and susceptibility to rapid tRNA decay by high-throughput in vivo analysis. Genes Dev. 28, 1721–1732 (2014).

29. Chen, X. et al. Visualizing RNA dynamics in live cells with bright and stable fluorescent RNAs. Nat. Biotechnol. 37, 1287–1293 (2019).

30. Andreasson, J. O. L., Savinov, A., Block, S. M. & Greenleaf, W. J. Comprehensive sequence-to-function mapping of cofactor-dependent RNA catalysis in the glmS ribozyme. Nat. Commun. 11, 1663 (2020).

31. Bai, J. et al. A protein-independent fluorescent RNA aptamer reporter system for plant genetic engineering. Nat. Commun. 11, 3847 (2020).

32. Torelli, E. et al. Light-Up Split Broccoli Aptamer as a Versatile Tool for RNA Assembly Monitoring in Cell-Free TX-TL Systems, Hybrid RNA/DNA Origami Tagging and DNA Biosensing. Int. J. Mol. Sci. 24, (2023).

33. Nilaratanakul, V., Hauer, D. A. & Griffin, D. E. Development of encoded Broccoli RNA aptamers for live cell imaging of alphavirus genomic and subgenomic RNAs. Sci. Rep. 10, 5233 (2020).

34. Filonov, G. S., Moon, J. D., Svensen, N. & Jaffrey, S. R. Broccoli: Rapid Selection of an RNA Mimic of Green Fluorescent Protein by Fluorescence-Based Selection and Directed Evolution. J. Am. Chem. Soc. 136, 16299–16308 (2014).

35. Kwok, C. K. & Merrick, C. J. G-Quadruplexes: Prediction, Characterization, and Biological Application. Trends Biotechnol. 35, 997–1013 (2017).

36. Hou, J., Guo, P., Wang, J., Han, D. & Tan, W. Artificial dynamic structure ensemble-guided rational design of a universal RNA aptamer-based sensing tag. Proc. Natl. Acad. Sci. U. S. A. 121, e2414793121 (2024).

37. Chen, Z. et al. Genetically encoded RNA-based sensors with Pepper fluorogenic aptamer. Nucleic Acids Res. 51, 8322–8336 (2023).

38. Huang, K. et al. Structure-based investigation of fluorogenic Pepper aptamer. Nat. Chem. Biol. 17, 1289–1295 (2021).

39. Chen, Z. et al. Genetically encoded RNA-based sensors with Pepper fluorogenic aptamer. Nucleic Acids Res. 51, 8322–8336 (2023).

40. Wang, Q. et al. Inert Pepper aptamer-mediated endogenous mRNA recognition and imaging in living cells. Nucleic Acids Res. 50, e84 (2022).

41. Tang, A. A. et al. Optimization of RNA Pepper Sensors for the Detection of Arbitrary RNA Targets. ACS Synth. Biol. 13, 498–508 (2024).

42. Rees, H. C., Gogacz, W., Li, N.-S., Koirala, D. P. & Piccirilli, J. A. Structural Basis for Fluorescence Activation by Pepper RNA. ACS Chem. Biol. (2022).

43. Johnson, S. R. et al. Computational scoring and experimental evaluation of enzymes generated by neural networks. Nat. Biotechnol. (2024) doi:10.1038/s41587-024-02214-2.

44. Shulgina, Y. et al. RNA language models predict mutations that improve RNA function. Preprint at 10.1101/2024.04.05.588317 (2024).

45. Hinton, G. E. Training Products of Experts by Minimizing Contrastive Divergence. Neural Comput. 14, 1771–1800 (2002).

46. PDF.

47. Domingo, J., Diss, G. & Lehner, B. Pairwise and higher-order genetic interactions during the evolution of a tRNA. Nature 558, 117–121 (2018).

48. Akiyama, M. & Sakakibara, Y. Informative RNA-base embedding for functional RNA structural alignment and clustering by deep representation learning. Preprint at 10.1101/2021.08.23.457433 (2021).

